# Axonal Transport Of An Insulin-Like Peptide Mrna Promotes Stress Recovery In *C. Elegans*

**DOI:** 10.1101/594689

**Authors:** Rashmi Chandra, Lisa Li, Zahabiya Husain, Shashwat Mishra, Joy Alcedo

## Abstract

Aberrations in insulin or insulin-like peptide (ILP) signaling in the brain causes many neurological diseases. Here we report that mRNAs of specific ILPs are surprisingly mobilized to the axons of *C. elegans* during stress. Transport of the ILP *ins-6* mRNA to axons facilitates recovery from stress, whereas loss of axonal mRNA delays recovery. In addition, the axonal traffic of *ins-6* mRNA is regulated by at least two opposing signals: one that depends on the insulin receptor DAF-2 and a kinesin-2 motor; and a second signal that is independent of DAF-2, but involves a kinesin-3 motor. While Golgi bodies that package nascent peptides, like ILPs, have not been previously found in *C. elegans* axons, we show that axons of stressed *C. elegans* have increased Golgi ready to package peptides for secretion. Thus, our findings present a mechanism that facilitates an animal’s rapid recovery from stress through axonal ILP mRNA mobilization.

## INTRODUCTION

The inability to recover from stress predisposes us to many neurological dysfunctions, such as post-traumatic brain injuries and neurodegeneration [reviewed in (*Blázquez et al., 2014; Esch et al., 2002; Frey, 2013; Koenen et al., 2008; Zeng et al., 2016*)]. Here we use the worm *C. elegans* to elucidate the mechanisms of stress adaptation and recovery, since the worm can switch its physiological state in response to stress [reviewed in (*Riddle and Albert, 1997*)]. Like in humans [(*Blessing et al., 2017; Marcovecchio and Chiarelli, 2012*); reviewed in (*Bedse et al., 2015; Blázquez et al., 2014; Boucher et al., 2014; Frey, 2013; Zeng et al., 2016*)], insulin-like peptides (ILPs) and their receptor enable the worms to endure and recover from stress (*Kimura et al., 1997; Riddle and Albert, 1997*).

In *C. elegans*, environmental stressors, like food scarcity and high population density, cause the first larval (L1) stage of reproductive growth to form dauers, the stress-induced arrested alternative to the third-stage larva (L3) of the growth program (*Golden and Riddle, 1984*). Compared to an L3, a dauer is more stress-resistant (*Riddle and Albert, 1997*). When dauers sense the return of favorable environments, they exit into the last larval (L4) stage and become a reproductive adult (*Golden and Riddle, 1984; Riddle et al., 1981*). These developmental switches are controlled by the combinatorial activities of specific ILPs from specific neurons (*Bargmann and Horvitz, 1991; Cornils et al., 2011; Fernandes de Abreu et al., 2014; Li et al., 2003; Pierce et al., 2001; Schackwitz et al., 1996*).

The *C. elegans* ILPs exist as a large family of peptides with 40 members (*Li et al., 2003; Pierce et al., 2001*), which are organized into a network where one ILP regulates the expression of other ILPs (*Fernandes de Abreu et al., 2014*). This inter-ILP network coordinates distinct subsets of ILPs that either regulate entry into or exit from dauer arrest (*Cornils et al., 2011; Fernandes de Abreu et al., 2014*). A major node of the network is the ILP *ins-6*, because *ins-6* expression is affected by the highest number of ILPs and *ins-6* in turn affects the expression of many other ILPs (*Fernandes de Abreu et al., 2014*). This is consistent with the highly pleiotropic phenotype of *ins-6*, which is also a major regulator of stress recovery (*Chen et al., 2013; Cornils et al., 2011; Fernandes de Abreu et al., 2014*).

Previously, we have shown that *ins-6* acts from ASJ sensory neurons to promote exit from the dauer state (*Cornils et al., 2011*). In this study, we have discovered that upon dauer arrest, *ins-6* mRNA is surprisingly transported to ASJ axons. Because we observe axonal *ins-6* mRNA only in dauers, we have investigated the significance and mechanism of axonal ILP mRNA transport in response to stress. We show that mRNAs of ILPs that promote dauer exit are trafficked to axons and that axonal *ins-6* mRNA promotes rapid dauer recovery. This is further supported by our finding that dauers also have enhanced axonal mobilization of Golgi bodies that can locally package ILP mRNA products for prompt secretion from axonal compartments.

Insulin signaling is conserved between worms and humans [reviewed in (*Alcedo and Zhang, 2013*)]. Stress also influences insulin, insulin growth factor and ILP relaxin mRNAs that are expressed in the human brain [reviewed in (*Blázquez et al., 2014; Fernandez and Torres-Aleman, 2012; Wilkinson et al., 2005*)]. Thus, our study raises the possibility that stress-induced axonal transport of ILP mRNAs and Golgi bodies in the worms is also present in humans to facilitate stress recovery.

## RESULTS

### The insulin-like peptide *ins-6* mRNA is trafficked to dauer axons

The decision between reproductive growth and stress-induced dauer arrest is regulated by specific sensory neurons, many of which are found in the amphid organ [*Figure S1A*; (*White et al., 1986*)]. For example, the amphid chemosensory neurons ASI and ADF act redundantly to inhibit entry into dauer arrest under non-stress environments (*Bargmann and Horvitz, 1991*). In contrast, the amphid chemosensory neuron ASJ has two opposing functions—it promotes dauer entry in response to stress and it promotes dauer exit under improved environments (*Bargmann and Horvitz, 1991; Schackwitz et al., 1996*). These neurons send their axons to a structure called the nerve ring (NR) bundle (*Figure S1A*), where the axons synapse to each other (*White et al., 1986*).

These amphid neurons express ILPs that regulate the switches between reproductive growth and dauer arrest (*Cornils et al., 2011; Fernandes de Abreu et al., 2014; Li et al., 2003; Pierce et al., 2001*). One of these ILPs is *ins-6*, which has a minor role in inhibiting dauer entry and a major role in facilitating exit from dauer after environments improve (*Cornils et al., 2011; Fernandes de Abreu et al., 2014*). Consistent with its function, we found through fluorescent *in situ* hybridization (FISH) that during the dauer entry decision stage (L1), endogenous *ins-6* mRNA was in the ASI neuronal soma, where it remained expressed in later stages (*Figures 1, S1B-S1H*). In a few second-stage larvae (L2s), *ins-6* also started to be expressed in the soma of the neuron ASJ (*Figure 1B, 1B’, 1E-1F*), where *ins-6* persisted as the animal developed into L3 and L4 (*Figures 1C-1D, 1C’-1D’, 1E-1F, S1D-S1H*).

**Fig 1.**
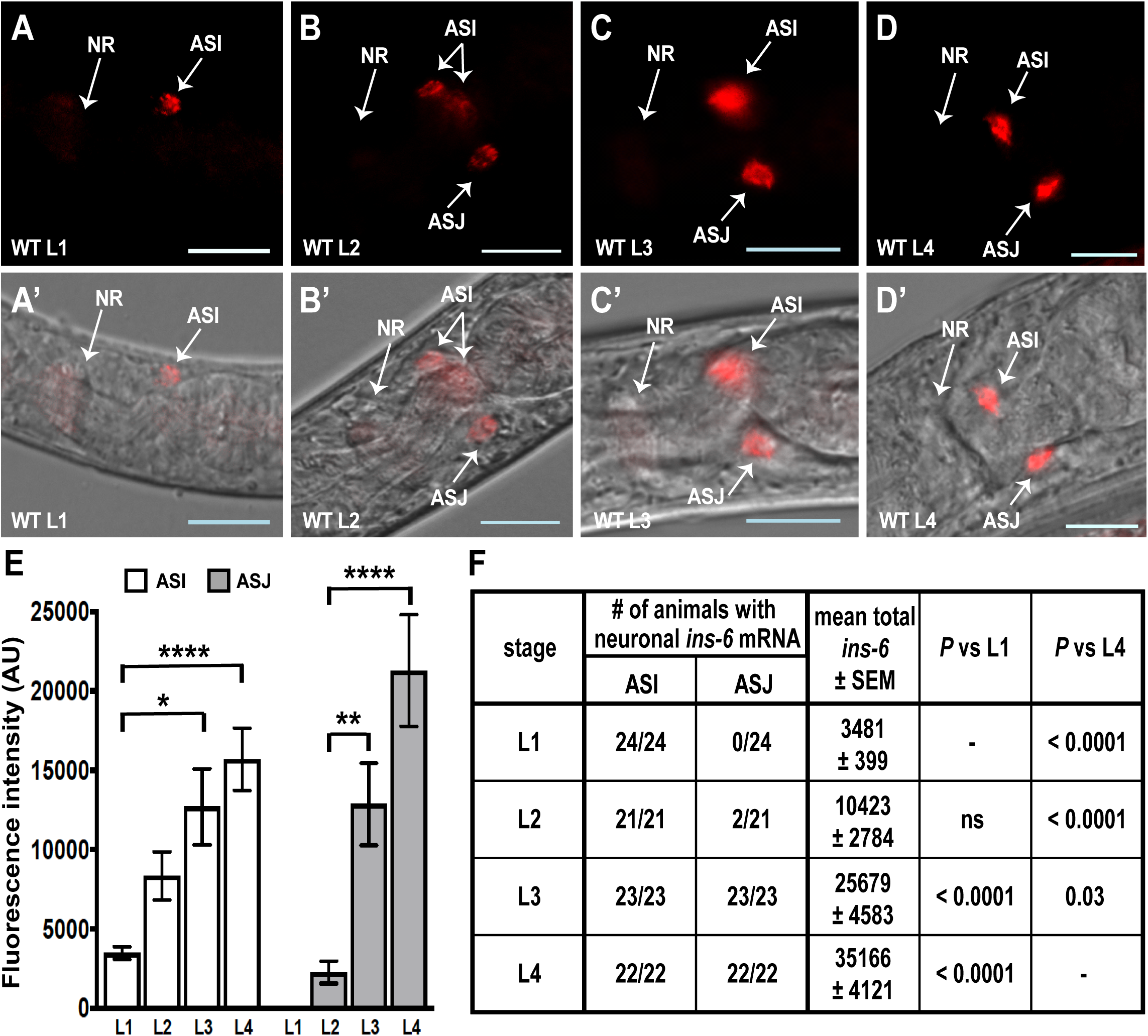
The ILP *ins-6* mRNA is expressed in the neuronal somas of wild-type larvae of the reproductive growth program. (A-D) show representative FISH images of *ins-6* mRNA in the somas of ASI and/or ASJ, while (A’-D’) show fluorescence overlay on corresponding DIC images of L1 (A and A’), L2 (B and B’), L3 (C and C’) and L4 (D and D’) larvae at 20°C. In this figure and later figures, anterior is to the left of each panel. Lateral views are shown, unless otherwise indicated. Scale bar is 10 μm. (E) Mean fluorescence intensities (± SEM) of *ins-6* mRNA in ASI versus ASJ are from 2 trials. (F) Distribution of larvae that express *ins-6* mRNA in ASI and/or ASJ and the mean total intensities (± SEM) of *ins-6* mRNA fluorescence from 2 trials. Statistical significance is determined by two-way ANOVA and Bonferroni correction. WT, wild-type; NR, nerve ring; AU, arbitrary units; ns, not significant; *, *P* < 0.05; **, *P* < 0.01; ****, *P* < 0.0001.

However, in dauers induced by high population density and starvation, *ins-6* mRNA was lost in ASI, but found in the ASJ soma and intriguingly in the NR axon bundle (*Figures 2A, 2B, 2A’, 2B’, 2E, S2A*). To confirm that the NR signal was specific to *ins-6* mRNA, we performed FISH on *ins-6* deletion mutant dauers and found that the *ins-6* NR signal was reduced to background in these mutants (*Figures 2C, 2C’, 2E, S2A*). We showed that *ins-6* mRNA in the NR was transported from the ASJ soma, since dauers that had genetically-ablated ASJ neurons also had background levels of the *ins-6* mRNA signal in the NR (*Figures 2D, 2D’, 2E, S2A*). The presence of *ins-6* mRNA in the ASJ axonal tract further supported the idea that *ins-6* mRNA was trafficked from the soma (*Figure 2F, 2G*). While the NR signal of *ins-6* mRNA was diffuse and broader than expected (*Figure 2A-2B, 2A’-2B’*), this signal resembled the broad signal, after the FISH protocol, of a GFP protein (*Figure 2H, 2H’*) that is expressed in only two pairs of axons (*Li et al., 2003*). We found that paraformaldehyde fixation and treatment with the hybridization buffer damaged the NR axonal fibers (*Figure 2I, 2I’*), which likely caused the scattering of the GFP protein or the fluorescent-labeled mRNAs from the axonal sources to the neighboring axons of the NR bundle.

**Fig 2.**
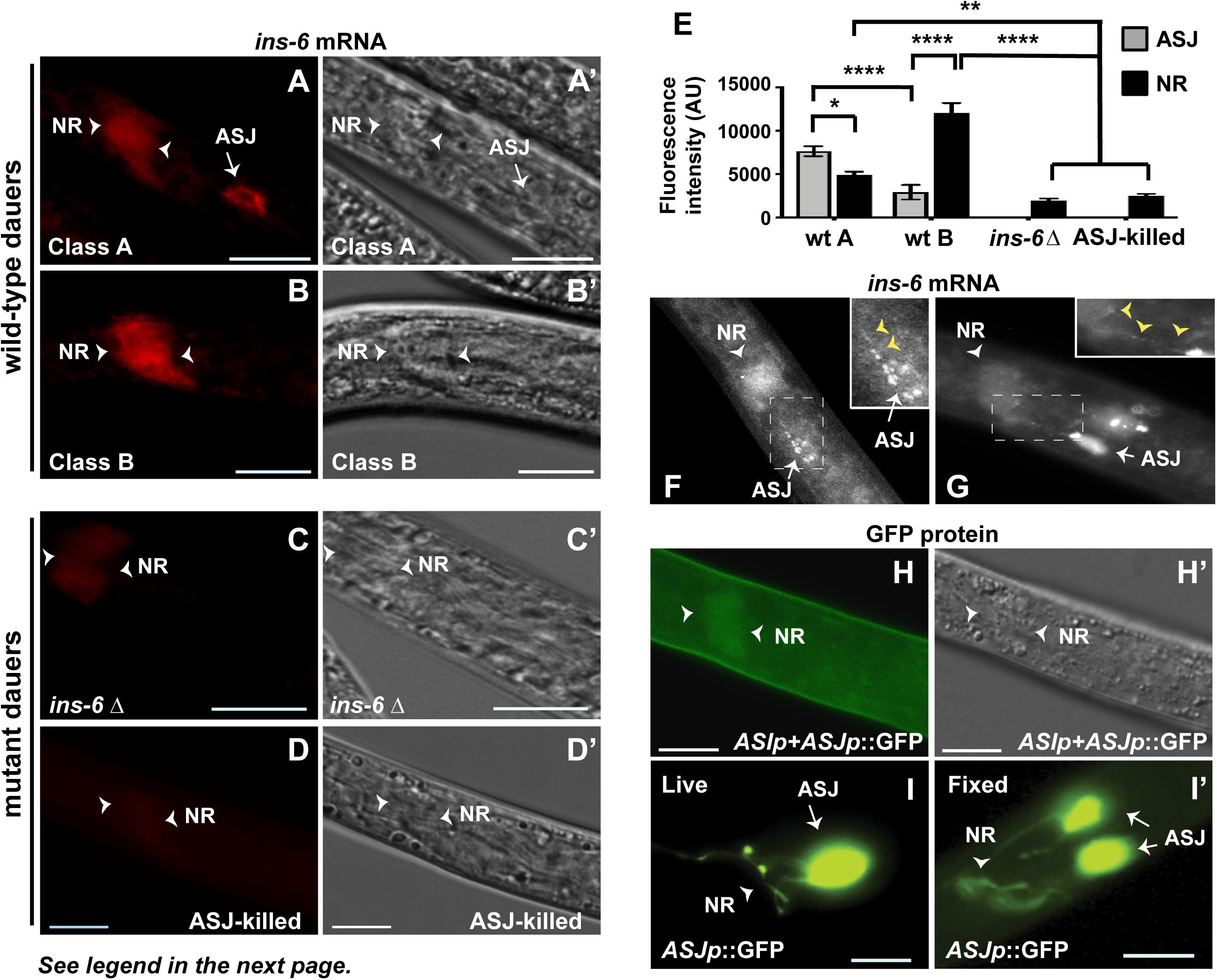
*ins-6* mRNA is transported to the NR from ASJ in dauers. (A-B) are representative fluorescent images of *ins-6* mRNA in the ASJ soma and/or NR of 5-day old dauers at 20°C; (A’-B’) are the corresponding DIC images of A and B. (A-B and A’-B’) are two classes of wild-type (wt) dauers based on *ins-6* mRNA levels in axons vs soma. (C-D) show *ins-6* mRNA fluorescent signals in mutant 5-day old dauers at 20°C; (C’-D’) are the corresponding DIC images of C and D. The *ins-6* mRNA signal is reduced to background in an *ins-6*(*tm2416)* deletion (Δ) mutant (C and C’) and in a dauer that has ASJ-killed neurons (D and D’). (E) Mean fluorescence intensities (± SEM) of *ins-6* mRNA in ASJ soma (grey bar) vs NR (black bar) from 3 trials. (F-G) FISH of *ins-6* mRNA in wild-type (F, lateral view) and *ins-6*-overexpressing (G, ventrolateral view) dauers. Insets are magnified from dotted boxes in main panels. Yellow arrowheads in insets point to the mRNA signal in ASJ axons. (H) GFP protein fluorescence (green) in a dauer that expresses GFP in two pairs of neurons, ASI and ASJ. (H’) Corresponding DIC image of H. (I) Image of a live, untreated dauer that expresses GFP protein from the ASJ neuron pair. (I’) Ventral view of a fixed and hybridization buffer-treated dauer that expresses GFP from the ASJ neuron pair, similar to I. Note the diffuse and broader GFP protein fluorescence from the NR (nerve ring) in this panel compared to the NR fluorescence in I. White arrowheads in A-D, A’-D’, H-I, H’-I’ point to the NR and/or its boundaries. AU, arbitrary units; ns, not significant; *, *P* < 0.05; **, *P* < 0.01; ****, *P* < 0.0001. Scale bar, 10 μm.

Wild-type dauers can be divided into two classes: class A represents dauers that have higher *ins-6* mRNA in the cell body, whereas class B are dauers that have higher *ins-6* in the axons (*Figures 2A, 2B, 2E, S2A*). Because starvation-induced dauers had *ins-6* mRNA in the NR (*Figure 2*), we then tested whether starvation alone was sufficient to traffic *ins-6* mRNA. While starvation reduced endogenous *ins-6* mRNA in both ASI and ASJ neurons, none of the starved larvae showed any *ins-6* mRNA in the NR (*Figure S2B-S2E*). Thus, the switch to the dauer state is required to traffic *ins-6* mRNA to axons. Once dauers exit to the last larval stage (post-dauer [PD] L4s), *ins-6* mRNA was also lost in the NR, but found in the ASJ soma of all animals and in the ASI soma of some animals (*Figure S1G-S1H*). Together our data suggest that axonal *ins-6* mRNA transport occurs only under certain stressed conditions, such as dauers.

### Dauer-exit promoting ILP mRNAs are in axons

Next, we asked whether all dauer-specific mRNAs are trafficked to the NR. We determined the dauer-dependent mRNA localization for other ILPs that regulate the switches between reproductive growth and dauer arrest (*Cornils et al., 2011; Fernandes de Abreu et al., 2014*). Like *ins-6*, the ILP *daf-28* inhibits dauer entry and promotes dauer recovery (*Cornils et al., 2011; Fernandes de Abreu et al., 2014; Kao et al., 2007; Li et al., 2003*). Similar to *ins-6*, we found that *daf-28* mRNA was present in the axons of wild-type dauers (*Figure 3B, 3B’*) and absent in *daf-28* deletion mutant dauers (*Figure 3C, 3C’*). Again, wild-type dauers can be divided into different classes based on the subcellular localization of *daf-28* mRNA. Class A dauers showed more *daf-28* mRNA in the somas of ASI and ASJ neurons (*Figure 3A, 3A’*); class B dauers showed more *daf-28* mRNA in the NR axon bundle (*Figure 3B, 3B’*). However, unlike class B *ins-6*-expressing animals, the class B *daf-28*-expressing animals were enriched in wild-type young dauers (2-day old; *Figure 3D*). About 78% of young dauers already had high levels of *daf-28* mRNA in the NR, whereas only 36% of this population had high levels of NR *ins-6* mRNA (*Figure 3D, 3E*). On the other hand, older wild-type dauers (5-day old) had decreased class B *daf-28*-expressing animals and increased class B *ins-6*-expressing animals (*Figure 3D*). Together these data show that both *daf-28* and *ins-6* mRNAs are trafficked to the dauer NR, but at different time points.

**Fig 3.**
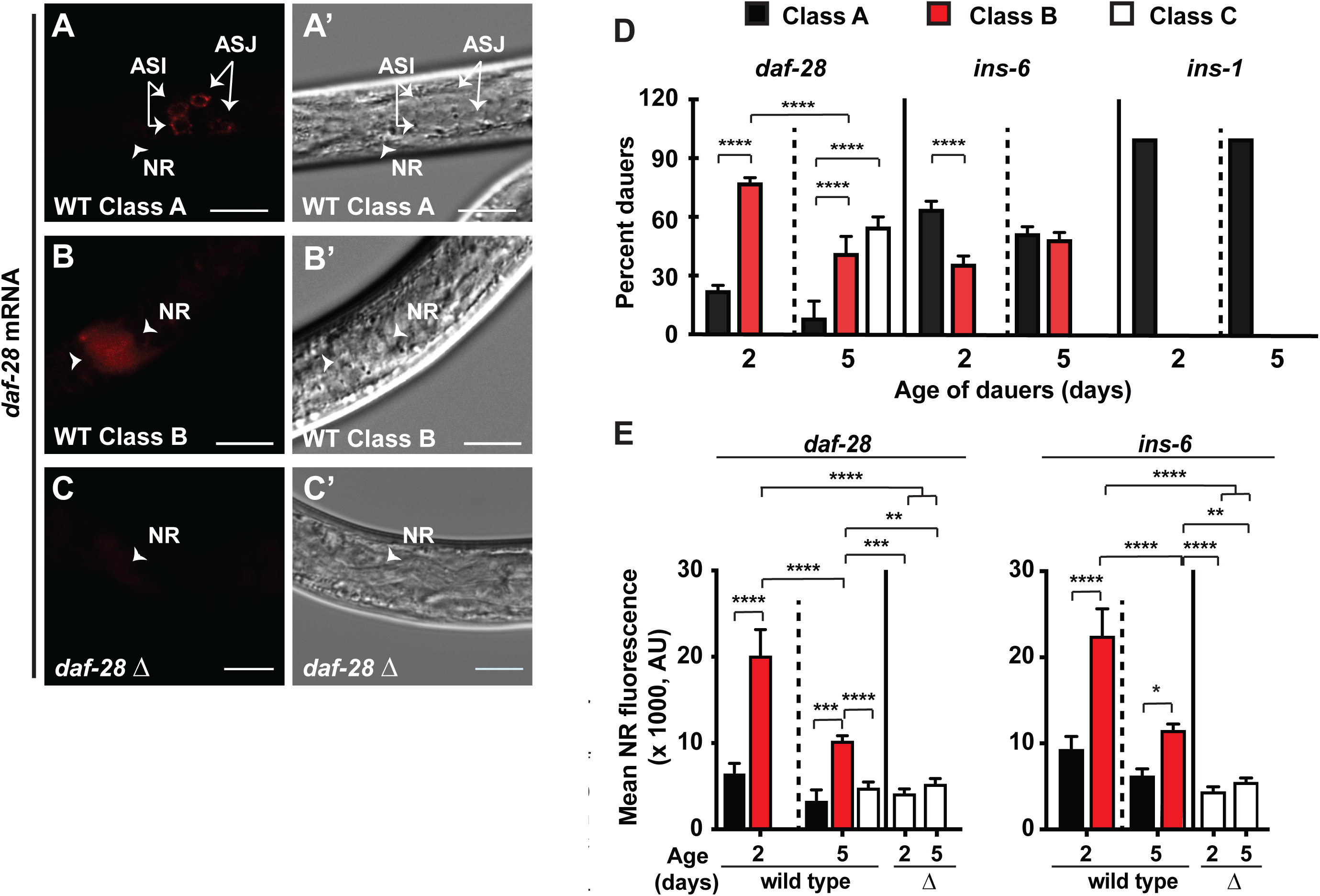
Exit-promoting ILP mRNAs are trafficked to dauer axons. (A-C) Fluorescent images of *daf-28* mRNA in ASI and ASJ somas or NR. (A’-C’) Corresponding DIC images at 20°C of 2-day (d) old WT (A-B) or *daf-28*(*tm2308)* deletion (Δ) mutant (C) dauers. (A-B and A’-B’) Two classes of WT dauers based on *daf-28* mRNA levels in axons vs somas. White arrowheads in A-C and A’-C’ point to the NR and/or its boundaries. Scale bar, 10 μm. (D) Mean distribution (± SEM) of 3 classes of 2-or 5-d old WT dauers at 20°C, based on specific ILP mRNA levels in somas vs NR. Class A dauers have more ILP mRNA in somas; class B, more ILP mRNA in NR; and class C, no expression in somas and NR. Number of WT dauers assayed: for *daf-28* mRNA, 24 (2-d) and 26 (5-d); for *ins-6*, 28 (2-d) and 30 (5-d); and for *ins-1*, 30 (2-d) and 36 (5-d), from at least 2 trials. (E) Mean NR signals (± SEM) of *daf-28* or *ins-6* mRNA in WT (from D) or the corresponding ILP deletion (Δ) mutant dauers. Number of *daf-28*(Δ) mutants assayed for *daf-28* mRNA are 26 (2-d) and 25 (5-d); and *ins-6*(Δ) mutants for *ins-6* mRNA, 25 (2-d) and 30 (5-d), from at least 2 trials. AU, arbitrary units. *, *P* < 0.05; **, *P* < 0.01; ***, *P* < 0.001; ****, *P* < 0.0001.

Unlike the NR axon bundle of young wild-type dauers, older dauers also had significantly less *daf-28* or *ins-6* mRNA in these axons (*Figure 3E*). In older wild-type animals, we observed the appearance of a third class of dauers in the population: the class C dauers that lack *daf-28* mRNA (*Figure 3D-3E*), which indicates that some ILP mRNAs are progressively lost in older dauers. Yet, the longer perdurance of *ins-6* mRNA than *daf-28* mRNA in the dauer NR (*Figure 3E*) highlights a more significant role for *ins-6* mRNA than *daf-28* in these axons. We previously showed that *ins-6* has a more important role in dauer exit than *daf-28* (*Cornils et al., 2011*), which suggests that the axonal transport of *ins-6* and *daf-28* mRNAs is to expedite dauer exit.

In support of this hypothesis, we found that the mRNAs of proteins that do not promote dauer exit were not trafficked to the NR [*Figures 3D, S3*; (*Cornils et al., 2011*)]. The mRNA of the ILP *ins-1*, which inhibits dauer exit (*Cornils et al., 2011*), was either expressed in the somas of multiple neurons (Class A1; *Figure S3C*) or in the soma of a single neuron in wild-type dauers (Class A2; *Figure S3D*). While the number of neuronal somas that express *ins-1* decreased in older dauers (*Figure S3A*), we did not detect *ins-1* mRNA in any dauer NR (*Figures 3D, S3B-S3E*).

Using animals that express GFP in the dauer ASI and ASJ neurons (*Li et al., 2003*), we also determined the location of *gfp* mRNA in these neurons. Since GFP plays no role in dauer exit, our hypothesis above predicts the absence of *gfp* mRNA in the dauer NR. Indeed, we observed no *gfp* mRNA in the NR, despite the presence of the GFP protein in these processes (*Figure S3F-S3F’*; see also *Figure 6C, 6C’*). Thus, lack of axonal *ins-1* and *gfp* mRNAs and presence of dauer-exit promoting ILP mRNAs in the NR suggest that axonal mRNA transport is required for dauer recovery.

### Insulin signaling and specific kinesins regulate axonal *ins-6* mRNA transport

To test our hypothesis, we asked how axonal trafficking of *ins-6* mRNA is regulated, which should also allow us to manipulate axonal *ins-6* mRNA levels and assess their effects on dauer recovery. The ILP receptor DAF-2 was previously shown to promote dauer exit (*Gems et al., 1998*). At 25°C, the *daf-2*(*e1368)* mutation leads to formation of dauers that exit after a few days, while the *daf-2*(*e1370)* mutation is a stronger allele that produces constitutive dauer arrest (*Gems et al., 1998*). We analyzed the axonal *ins-6* mRNA levels in dauers of both *daf-2* alleles at 25°C. We focused on 3-day old dauers, since wild type at this age at 25°C showed a distribution of class A and class B *ins-6*-expressing dauers comparable to 5-day old wild type at 20°C (*Figure 4*). Compared to wild-type dauers, we found that both *daf-2* mutants had much fewer class B dauers (*Figures 4A, S4A*). The lack of exit in *daf-2*(*e1370)* dauers at 25°C (*Gems et al., 1998*) is likely due to loss of *ins-6* mRNA in some of these animals, since we also observed the emergence of class C dauers that had no *ins-6* expression in the ASJ soma and NR (*Figure 4A*). Thus, the wild-type DAF-2 receptor stimulates *ins-6* mRNA transport to the NR to induce dauer recovery.

**Fig 4.**
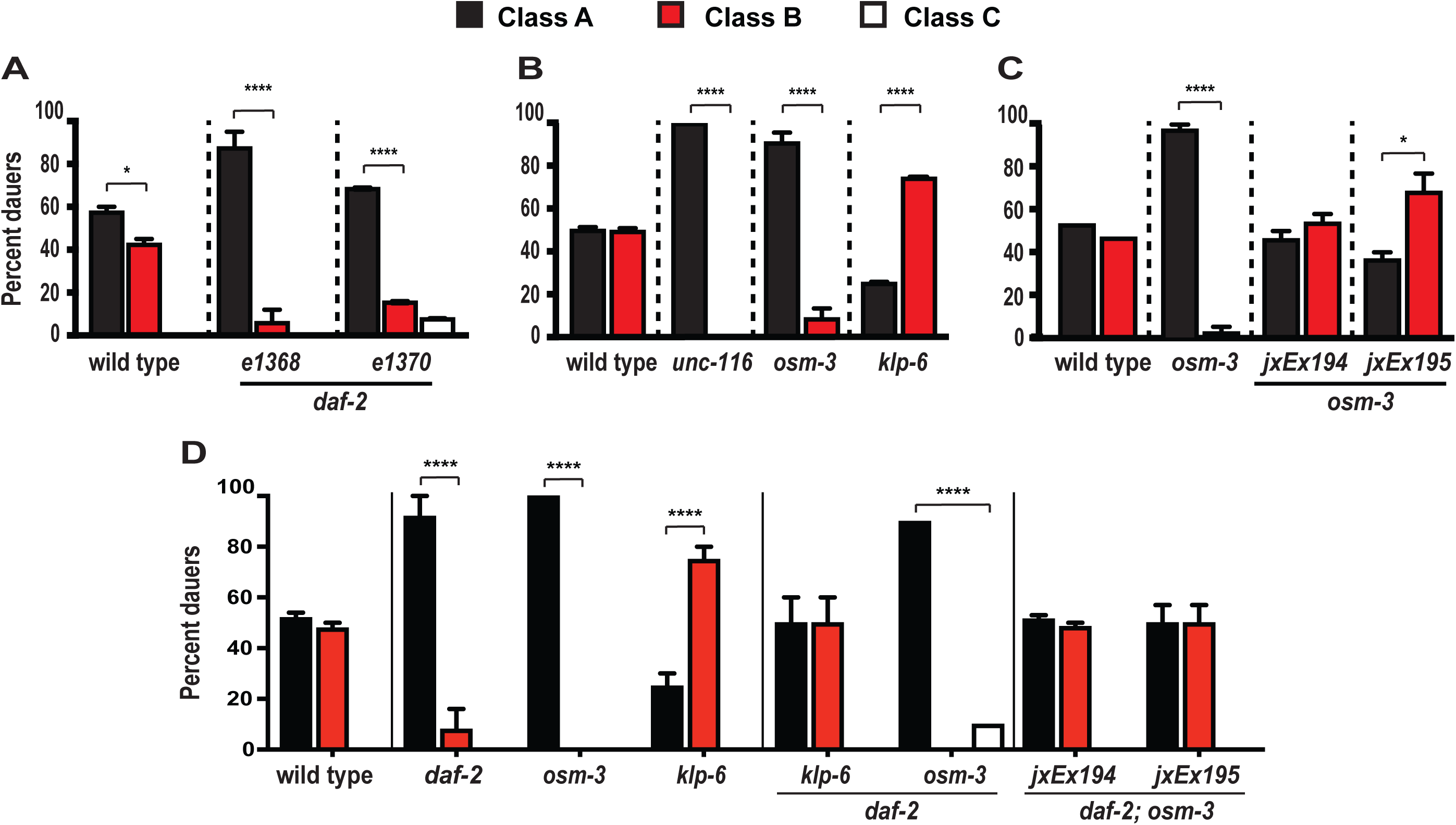
Insulin signaling and specific kinesins modulate axonal *ins-6* mRNA levels. (A-D) Mean distribution (± SEM) of three classes of wild-type or mutant dauers. In this figure and subsequent figures, class A dauers have more *ins-6* mRNA in the ASJ soma; class B, more *ins-6* mRNA in the NR; and class C, no *ins-6* expression in somas and NR. (A) The 3-day old dauers of *daf-2* alleles, *e1368* and *e1370*, compared to 3-day old wild-type dauers at 25°C. (B) Different kinesin mutations, *unc-116*(*e2310), osm-3*(*p802)*, and *klp-6*(*sy511)*, have different effects on the axonal *ins-6* mRNA of 5-day old dauers at 20°C. (C) ASJ-specific expression of *osm-3* (*jxEx194* or *jxEx195*) rescues axonal *ins-6* mRNA levels in 5-day old dauers at 20°C. (D) *osm-3*(*p802)* or *klp-6*(*sy511)* in the *daf-2*(*e1368)* mutant background alters the axonal *ins-6* mRNA phenotype of the *daf-2* mutation in 3-day old dauers at 25°C. ASJ-specific rescue of *osm-3* (*jxEx194* or *jxEx195*) in *daf-2*; *osm-3* double mutants restores class B dauers. Statistical comparisons within each strain of animals are shown. See *Figure S4* for statistical comparisons across different strains. *, *P* < 0.05; ****, *P* < 0.0001.

Axonal transport depends on kinesins, which are the motors that carry cargo along the microtubules that run the length of an axon [reviewed in (*Scholey, 2013; Siddiqui, 2002*)]. *C. elegans* has multiple kinesins, some of which are expressed in neurons (*Scholey, 2013; Siddiqui, 2002*). Two of these kinesins represent members of the kinesin-1 and kinesin-2 families, which have been implicated in transporting mRNAs to different subcellular compartments in other animals (*Kanai et al., 2004; Messitt et al., 2008*). In *C. elegans, unc-116* is a kinesin-1 motor, *osm-3* is a kinesin-2 motor, and both kinesins are present in ASJ (*Sakamoto et al., 2005; Scholey, 2013; Tabish et al., 1995*). In *unc-116* or *osm-3* loss-of-function mutants, class B *ins-6-*expressing dauers are lost (*Figures 4B, S4B*), which indicates that wild-type *unc-116* and *osm-3* are required in transporting *ins-6* mRNA to dauer axons. The rescue of *osm-3* in ASJ alone, *jxEx194* and *jxEx195*, also rescued the class A and class B distributions of *ins-6*-expressing dauers, which shows that OSM-3 kinesin acts in ASJ to mobilize *ins-6* mRNA to axons (*Figures 4C, S4C*).

Next, we asked if neuronal kinesins that are absent from ASJ also exert a similar effect on *ins-6* mRNA, such as *klp-6*, which is a kinesin-3 motor (*Peden and Barr, 2005*). Interestingly, a partial loss-of-function mutation in *klp-6* (*Peden and Barr, 2005*) led to an enrichment in class B *ins-6*-expressing dauers (*Figures 4B, S4B*), which suggests that wild-type *klp-6* inhibits *ins-6* mRNA transport. Because KLP-6 is not in ASJ (*Peden and Barr, 2005*), this suggests that the *klp-6*-dependent regulation of axonal *ins-6* mRNA involves a signal secreted from *klp-6*-expressing cells. Considering the *daf-2*-dependence of *ins-6* mRNA transport (*Figures 4A, S4A*), this signal might be another ILP that is a DAF-2 receptor ligand.

Yet, the epistasis between *daf-2* and *klp-6* implies the involvement of more than one signal (*Figures 4D, S4D*). Similar to 20°C, *klp-6* single mutants again had more class B *ins-6*-expressing dauers at 25°C, unlike *daf-2* single mutant or wild-type dauers under the same conditions (*Figures 4D, S4D*). In contrast, the *daf-2 klp-6* double mutants exhibited the wild-type dauer distribution phenotype (*Figures 4D, S4D*). Since the *daf-2 klp-6* double mutants showed an intermediate phenotype between those of *daf-2* and *klp-6* single mutants (*Figures 4D, S4D*), this implicates a second signal that acts parallel to the DAF-2 ligand in the *klp-6*-dependent inhibition of axonal *ins-6* mRNA transport.

In comparison, the kinesin OSM-3 likely functions downstream of the DAF-2 receptor. As at 20°C, the severe *osm-3* single mutants again lost class B *ins-6*-expressing dauers at 25°C, which resembled *daf-2* single mutants (*Figures 4D, S4D*). Similar to the stronger *daf-2*(*e1370)* single mutant (*Figures 4A, S4A*), the *daf-2*(*e1368)*; *osm-3* double mutants produced class C dauers that lost all *ins-6* expression (*Figures 4D, S4D*). However, the ASJ-specific rescue of *osm-3* in the *daf-2*(*e1368)*; *osm-3* double mutants restored the wild-type class A and class B distributions of *ins-6*-expressing dauers (*Figures 4D, S4D*). This suggests that OSM-3 activity in ASJ is downstream of DAF-2 in promoting *ins-6* mRNA transport to dauer axons.

### Axonal *ins-6* mRNA levels modulate dauer recovery

The altered *ins-6* mRNA distribution in the different mutants allowed us to test the significance of axonal *ins-6* mRNA on dauer recovery. Because *daf-2*(*e1368)* mutant dauers exit after a few days (*Gems et al., 1998*), we analyzed the dauer recovery of *daf-2*(*e1368)* in the presence versus absence of the wild-type kinesins that regulate axonal *ins-6* mRNA levels. The mutations in kinesins *osm-3* and *unc-116*, which failed to transport *ins-6* mRNA axonally (*Figures 4B-4D, S4*), significantly delayed the exit of *daf-2*(*e1368)* dauers (*Figure 5A, Table S1*). The ASJ-specific rescue of *osm-3*, which restored the high axonal *ins-6*-expressing class B dauers (*Figures 4D, S4*), also rescued the delayed dauer exit of *daf-2*; *osm-3* double mutants back to the exit phenotype of *daf-2* single mutants (*Figure 5C, Table S1*). These suggest that axonal *ins-6* mRNA expedites dauer recovery.

**Fig 5.**
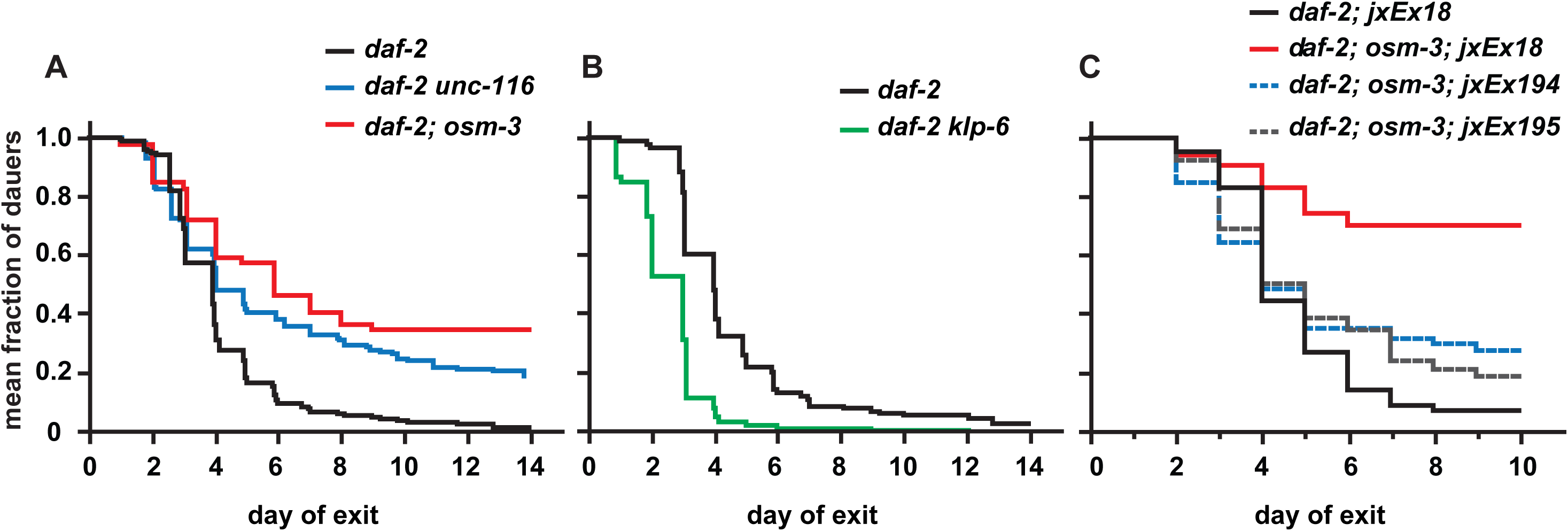
Axonal *ins-6* mRNA facilitates stress recovery. (A-C) The dauer exit rates at 25°C of animals carrying different kinesin mutations, *unc-116*(*e2310), osm-3*(*p802)* or *klp-6*(*sy511)*, in the *daf-2*(*e1368)* mutant background. (A-B) Animals that carry a kinesin mutation have a dauer exit rate that is significantly different from the *daf-2*(*e1368)* control by *P* < 0.0001, according to the logrank test. Each curve represents cumulative data from 3-5 trials. (C) ASJ-specific rescue of *osm-3* (*jxEx194* or *jxEx195*) in *daf-2; osm-3* double mutants restores dauer exit rates back to the *daf-2*(*e1368)* control, which carries the *ofm-1::gfp* coinjection marker (*jxEx18*) that is also present in *jxEx194* or *jxEx195*. Each curve represents cumulative data from 2 trials. The complete statistical comparisons between the dauer exit phenotypes of the different groups of animals are shown in *Table S1*.

**Fig 6.**
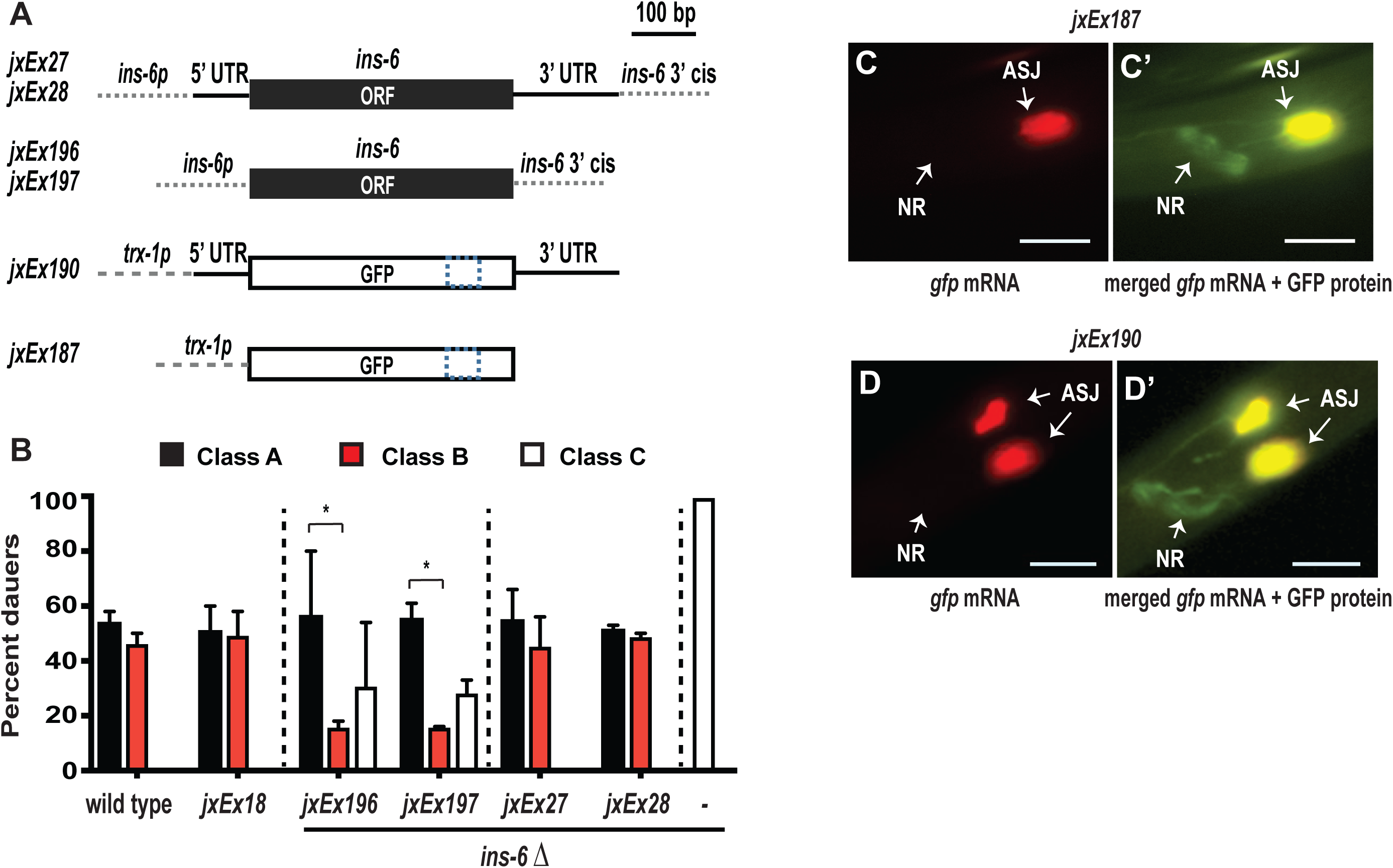
*ins-6* UTR sequences are partly necessary, but insufficient to transport *ins-6* mRNA axonally. (A) Diagrams of transgenes used. *jxEx27* or *jxEx28* is an *ins-6* rescuing transgene with the full *ins-6* genomic locus (*Cornils et al., 2011*); *jxEx196* or *jxEx197*, the *ins-6* rescuing transgene lacking UTRs. *jxEx190* is a GFP cDNA flanked by the *ins-6* UTRs and specifi-cally expressed in ASJ. *jxEx187* is an ASJ-expressed GFP construct that lacks the *ins-6* UTRs. The dotted box in GFP denotes its greater length than the *ins-6* ORF. The dotted lines of the *ins-6* or *trx-1* promoter or *ins-6* 3’ *cis* regulatory region indicate that they are not drawn to scale. (B) Mean distribution (± SEM) at 20°C of three classes of wild-type or *ins-6* mutant 5-day old dauers that carry the indicated transgenes. *jxEx18* depicts the *ofm-1::gfp* coinjection marker, which has no effect on the class distribution of dauers. The *ins-6* transgenes (*jxEx27, jxEx28, jxEx196* or *jxEx197*) were injected with the *ofm-1::gfp* coinjection marker into the *ins-6*(*tm2416)* deletion (Δ) mutant. Number of dauers assayed were 20-30 per strain. *, *P* < 0.05. (C-D) *gfp* mRNA fluorescence in a 5-day old dauer that carries *jxEx187* (C) or *jxEx190* (D) at 20°C. (C’-D’) Merged mRNA (red) and protein (green) fluorescence of GFP in 5-day old dauers that carry the indicated transgenes. Yellow indicates colocalization of red and green fluorescence. Scale bar, 10 μm.

Indeed, loss of *klp-6*, which enriched for the high axonal *ins-6*-expressing dauer population (*Figures 4B, 4D, S4*), caused *daf-2*(*e1368)* mutant dauers to exit earlier to L4 (*Figure 5B, Table S1*). Interestingly, both the *daf-2*; *osm-3* double mutants that have wild-type *osm-3* in ASJ and the *daf-2 klp-6* double mutants showed similar distributions of class A and class B *ins-6*-expressing dauers (*Figures 4D, S4*), but different dauer exit phenotypes compared to *daf-2* single mutants (*Figure 5B-5C, Table S1*). This implies that the presence of axonal *ins-6* mRNA in dauers is only one of the requirements needed to induce exit. However, because restoring class B *ins-6*-expressing dauers to a population accelerated dauer exit [compare (i) *daf-2*; *osm-3*; *jxEx194* or *jxEx195* transgenic animals to *daf-2*; *osm-3* non-rescued double mutants or (ii) *daf-2 klp-6* double mutants to *daf-2* single mutants], this highlights the importance of axonal *ins-6* mRNA in dauer recovery.

### Untranslated regions of *ins-6* are partly necessary for its mRNA transport

Untranslated regions (UTRs) control gene expression [reviewed in (*Decker and Parker, 1995*)] through mRNA stability (*Brown et al., 1993; Scheper et al., 1995*) and/or subcellular localization (*Bertrand et al., 1998; Gunkel et al., 1998; Thio et al., 2000*). Since *ins-6* mRNA has both 5’ and 3’ UTRs (*Figure 6A*), we asked if axonal trafficking of *ins-6* mRNA requires its UTRs. We generated constructs that lacked both 5’ and 3’ UTRs of *ins-6* (*Figure 6A*) and compared their mRNA localization, *jxEx196* and *jxEx197*, with the mRNA localization of constructs that carry the whole *ins-6* genomic locus, *jxEx27* and *jxEx28* (*Figure 6A*), in the *ins-6* deletion background (*Figure 6B*). The full *ins-6* genomic locus, *jxEx27* or *jxEx28*, rescued the *ins-6* deletion phenotype back to wild type (*Figure 6B*). In contrast, the UTR-less constructs, *jxEx196* or *jxEx197*, only partly rescued the *ins-6* mutant phenotype (*Figure 6B*), which show that UTRs modulate *ins-6* mRNA levels. Since these animals also exhibited class B dauers (*Figure 6B*), then loss of UTRs still allows *ins-6* mRNA transport to axons. However, the UTR-less animals had significantly fewer class B dauers than the full-rescue *ins-6* lines (*Figure 6B*), which means that UTRs are at least partly necessary for axonal *ins-6* mRNA.

Next, we asked if *ins-6* UTRs are sufficient to carry any mRNA to dauer axons. *gfp* mRNA alone was not trafficked to dauer axons, whether *gfp* was expressed from ASI and ASJ (*Figure S3F-S3F’*) or from ASJ only (*jxEx187*; *Figure 6A, 6C, 6C’*). We tagged *gfp* mRNA with the *ins-6* 5’ and 3’ UTRs and expressed it from ASJ (*jxEx190*; *Figure 6A*). Although the GFP protein can be found in the dauer axons, the *gfp*-*ins-6* UTR mRNA hybrid remained in the ASJ soma (*Figure 6D, 6D’*). Thus, while *ins-6* UTRs are partly necessary to traffic *ins-6* mRNA to axons, the UTRs are insufficient for axonal mRNA transport.

### Stress enhances Golgi mobilization to axons

Like other secreted proteins, ILP mRNAs need to be processed and packaged for secretion, but previous literature only reported the presence of rough endoplasmic reticulum in wild-type *C. elegans* axons (*Edwards et al., 2013*). We determined if *C. elegans* axons also contain Golgi that would be needed to package locally translated peptides. We used a pan-neuronally expressed YFP-tagged Golgi marker, α-mannosidase II [*aman-2*; (*Edwards et al., 2013; Sumakovic et al., 2009*)]. *C. elegans* larvae, like dauer and its alternative form, L3, have Golgi in neuronal somas, dendrites, and axons, which include the dorsal cord axons and NR, but not the commissures (*Figure 7, Videos 1-2*). Notably, dauers had more axonal Golgi (*Figure 7E, Video 2*), which suggests that stress increases Golgi mobilization to process newly synthesized proteins in axons.

**Fig 7.**
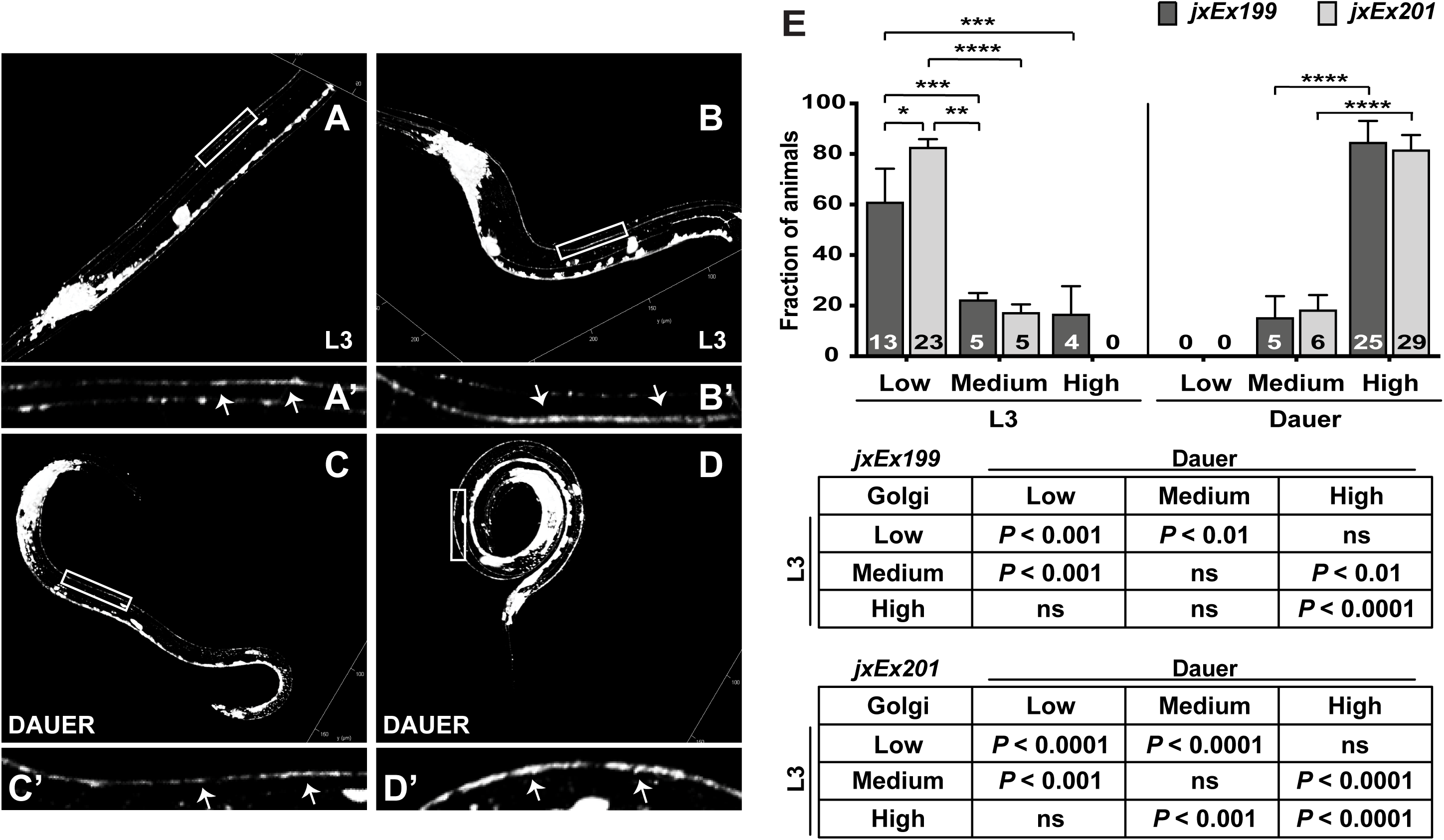
Stress enhances Golgi mobilization to the dorsal cord (DC) axons. (A-D) 3D-reconstruction of *jxEx199* larvae that express the Golgi marker AMAN-2::YFP (*Edwards et al., 2013; Sumakovic et al., 2009*) at 20°C. (A-B) L3 larvae with medium (A) or high (B) amounts of Golgi in DC axons. (A’-B’) Higher magnification of a 50-μm section of the L3 DC axons (indicated by arrows; boxes in A and B). (C-D) Dauers with medium (C) or high (D) amounts of Golgi in the DC axons. (C’-D’) Higher magnification of a 50-μm section of the dauer DC axons (indicated by arrows; boxes in C and D). (E) Mean fractions (± SEM) of L3s and dauers that show different levels of Golgi bodies in DC axons from 3 trials, using 2 lines that express AMAN-2::YFP, *jxEx199* and *jxEx201*. Tables show the statistical comparisons between L3s and dauers of each line.

## DISCUSSION

Aberrant insulin signaling hampers recovery from stress. Because insulin signaling ensures homeostasis (*Blázquez et al., 2014; Fernandes de Abreu et al., 2014*), ligands for this pathway undergo multiple levels of regulation to enable the pathway to function with high precision (*Fu et al., 2013*). While the significance of ILP mRNA subcellular localization has yet to be considered, we show that certain stressors localize mRNAs of specific ILPs through distinct kinesin activities to *C. elegans* axons, which is accompanied by an increase in axonal Golgi mobilization. Importantly, our study suggests that these ILP mRNAs must be in the axons and locally translated and packaged for prompt secretion to facilitate stress recovery. This illustrates a novel mechanism where ILP mRNAs maintain neuronal plasticity during stress.

### Certain ILP mRNAs are localized to axons by specific stressors to promote recovery

The environment alters *ins-6* expression at the transcriptional and post-transcriptional levels. *ins-6* is first expressed in one pair (ASI) and later in two pairs (ASI and ASJ) of sensory neurons under optimal environments (*Figures 1, S1*). Like other ILP mRNAs that are present in neuronal somas of *Drosophila* and vertebrate brains (*Banerjee et al., 2010; Bathgate et al., 2013; Brogiolo et al., 2001; Broughton et al., 2005; Ma et al., 2009*), *ins-6* mRNA is in ASI and ASJ somas under non-stress conditions. In starvation-induced dauers, *ins-6* is off in ASI, but remains expressed in ASJ. Importantly, *ins-6* mRNA is mobilized to dauer ASJ axons. This ILP mRNA localization presumably leads to a prompt resetting of homeostasis after a major environmental fluctuation.

Yet, why do we see axonal *ins-6* mRNA in dauers (*Figure 2*), but not in starved non-dauers (*Figure S2B-S2E*)? This is likely because the dauer itself is an adaptive response that the animal uses to survive a potentially protracted, harsh environmental change, as it awaits a better environment (*Riddle and Albert, 1997*). For this reason, the dauer program should include a priming mechanism that promotes recovery. The axonal transport of an ILP mRNA, whose peptide product is required for dauer exit, could serve as a priming mechanism. Because dauers have decreased global transcription (*Dalley and Golomb, 1992; Wang and Kim, 2003*), the local translation of a previously existing mRNA will lead to a faster response to improved environments. This is important, since a prolonged stay in the dauer state can lead to deficits in animals that eventually exit (*Kim and Paik, 2008*). Thus, it is not surprising that the mRNAs of ILPs that promote exit are the mRNAs that are localized to axons (*Figures 2, 3*), whereas mRNAs of proteins that do not promote exit (*Cornils et al., 2011*) are absent in axons (*Figure S3*).

The difference in temporal axonal mRNA localization patterns for *ins-6* and *daf-28* (*Figure 3D-3E*) hint at different exit requirements for young versus old dauers. However, the longer perdurance of axonal *ins-6* mRNA than *daf-28* mRNA (*Figure 3D-3E*) is consistent with the larger role of *ins-6* versus *daf-28* in promoting dauer recovery (*Cornils et al., 2011*). An increase in axonal *ins-6* mRNA in a dauer population enhances exit, whereas a decrease in axonal *ins-6* mRNA delays exit (*Figures 4, S4, 5, Table S1*). As an animal that has recovered from dauer also shows no axonal *ins-6* mRNA (*Figure S1G-S1H*), this indicates that *ins-6* mRNA is not needed in axons in post-recovery, further highlighting the significance of axonal ILP mRNA in stressed animals.

### Insulin signaling regulates the mRNA localization of at least one of its ligands

Insulin receptor activity promotes mRNA expression (*Fernandes de Abreu et al., 2014; Kaplan et al., 2019; Leibiger et al., 1998; Li et al., 2003; Murphy et al., 2003*), translation (*Lee et al., 2012*), and secretion of some of its ligands (*Kulkarni et al., 1999*) across species. Here we show that insulin receptor activity also promotes the mobilization of the mRNA of at least one of its ligands to specific subcellular compartments, *e.g.*, the *C. elegans* axons (*Figures 4, S4*). This positive feedback regulation in insulin signaling allows the rapid responses (*Kaplan et al., 2019*) that are necessary in switching from one state to another, such as exit from dauer to post-dauer state.

The *C. elegans* insulin receptor DAF-2 stimulates dauer exit (*Gems et al., 1998*), at least by regulating the fate of *ins-6* mRNA (*Figures 4, S4*). Our data suggest that DAF-2 signaling activates OSM-3 kinesin in ASJ to promote *ins-6* mRNA traffic to axons (*Figures 4, S4*), which facilitates dauer exit (*Figure 5B-5C, Table S1*). Intriguingly, animals that have the weak loss-of-function *daf-2*(*e1368)* and the strong *osm-3* loss of function also show the more severe phenotype of the strong *daf-2* loss-of-function mutant *e1370*, where some dauers have lost *ins-6* expression in both soma and axon (*Figures 4, S4*). This suggests that both DAF-2 and OSM-3 have two roles: one is to promote axonal *ins-6* mRNA transport and the other is to inhibit loss of *ins-6* mRNA. While this is similar to the function of *ins-6* UTRs, which are partly required for *ins-6* mRNA transport and steady-state levels (*Figure 6*), it remains to be seen if OSM-3 is the kinesin that mobilizes *ins-6* mRNA to axons via the *ins-6* UTRs in response to DAF-2 signaling.

It must be noted that axonal *ins-6* mRNA is only one of the requirements for dauer exit. Wild-type OSM-3 in ASJ rescues the severe delay in dauer exit of *daf-2*; *osm-3* double mutants back to the exit phenotype of *daf-2* single mutants (*Figure 5C, Table S1*), although the rescued double mutants have axonal *ins-6* mRNA brought back to wild-type levels (*Figures 4D, S4D*). Nonetheless, the importance of axonal *ins-6* mRNA in dauer recovery is evident in the enhanced exit of dauers (*Figure 5B-5C, Table S1*) to which axonal *ins-6* mRNA is restored (*Figures 4D, S4D*).

Our data also suggest that the *daf-2*-dependent axonal localization of *ins-6* mRNA in ASJ is antagonized by a *daf-2*-independent, but *klp-6*-dependent signal that is secreted from other neurons (*Figures 4D, S4D*). The regulation of axonal *ins-6* mRNA transport by these opposing signals underscores the idea that *ins-6* mRNA must be in axons only when needed, such as when the animal has to prepare for exit into a much better environment that supports survival.

### Transport of Golgi bodies to larval axons increases with stress

The mRNAs transported to axons are believed to be locally translated to expedite protein processing near the synaptic zone [reviewed in (*Holt and Schuman, 2013; Jansen, 2001*)], but an ILP peptide also needs to be packaged from Golgi prior to its secretion. In support of axonal translation, the rough endoplasmic reticulum has been found in axons of *C. elegans* and other animals [(*Edwards et al., 2013*); reviewed in (*Spaulding and Burgess, 2017*)]. In contrast, axonal Golgi has been rarely reported in other species (*Morest, 1971; Wigglesworth, 1960; Willis et al., 2011*) and has not been observed in adult *C. elegans* (*Edwards et al., 2013*). Interestingly, Golgi is in *Drosophila* larval axons (*Cao et al., 2014*), which suggests that *C. elegans* larvae will also have axonal Golgi. Indeed, we find Golgi in the axons of L3 and at much higher levels in the axons of dauers (*Figure 7, Videos 1-2*), which suggests that stress increases axonal Golgi mobilization.

Why should *C. elegans* larvae have axonal Golgi? Axonal translation is important in axon migration and synaptogenesis (*Holt and Schuman, 2013*). In *C. elegans*, synapse formation along the length of axons and axon outgrowth continue during larval stages [(*Gujar et al., 2017; Lipton et al., 2018; Shen and Bargmann, 2003; White et al., 1978; White et al., 1986; Zhao and Nonet, 2000*); reviewed in (*Chisholm et al., 2016*)]. These might require axonal Golgi to package the necessary proteins in larvae (*Colón-Ramos et al., 2007; Shen and Bargmann, 2003*).

The dauer axons can also be remodeled (*Schroeder et al., 2013*), which can lead to synaptogenesis. Since dauers have reduced global transcription (*Dalley and Golomb, 1992; Wang and Kim, 2003*), it is possible that a number of mRNAs, besides those involved in stress recovery, are axonally trafficked. The packaging of locally translated membrane or secreted proteins that promote axon outgrowths, synaptogenesis, and stress adaptation and recovery will require the presence of axonal Golgi. Thus, increased mobilization of Golgi and specific ILP mRNAs to dauer axons is a needed response in stress adaptation and recovery. This suggests a mechanism that promotes plasticity during stress and allows for optimal recovery from stress, a mechanism that might be conserved across metazoans.

## MATERIALS AND METHODS

### Experimental model

Wild-type (N2) or mutant *C. elegans* hermaphrodites were grown at 20°C or 25°C, as indicated, using standard protocols (*Brenner, 1974; Cornils et al., 2011*). Worms were grown on *E. coli* OP50 before each assay. Dauers were induced naturally through overcrowding and starvation (*Golden and Riddle, 1984*), except for the *daf-2* mutants that routinely form dauers in the presence of abundant food at 25°C (*Gems et al., 1998*). All worm mutants in this study were backcrossed at least four times to our lab N2 strain before any analysis was performed, including the ASJ-ablated animals that have the *trx-1p::ICE* transgene (*Cornils et al., 2011*) integrated into their genome (generous gift of Miriam Goodman).

#### Transgene constructions

To generate an ASJ-specific rescue of *osm-3*, we drove the expression of a GFP-tagged *osm-3* cDNA from an ASJ-specific promoter [pQZ93; (*Cornils et al., 2011*)]. We replaced the *srh-114* promoter of OSM-3::GFP in the pAGB060 plasmid (generous gift from Piali Sengupta) with the ASJ-specific *trx-1* promoter (*Cornils et al., 2011; Miranda-Vizuete et al., 2006*). To generate the UTR-less *ins-6* transgene (pQZ97), we deleted the 72-bp 5’ UTR and 110 bp from the 3’ UTR of the full *ins-6* rescuing construct pQZ11 (*Cornils et al., 2011*). The polyA signal was kept at the 3’ UTR of pQZ97. To create the ASJ-specific GFP-expressing lines with the *ins-6* 5’ and 3’ UTRs (pQZ91), we fused the 72-bp *ins-6* 5’ UTR to the 5’ end of the GFP cDNA in pQZ34, which used the *trx-1* promoter to drive GFP expression (*Figure 6A*) from the vector backbone of pPD95.77 (generous gift from Andrew Fire). We also inserted the 124-bp *ins-6* 3’ UTR to the 3’ end of the GFP cDNA in pQZ34 (*Figure 6A*).

#### Transgenic worms

To create the ASJ-specific *osm-3* rescue lines, we injected 25 ng/μl of pQZ93 with 25 ng/μl of the coinjection marker *ofm-1::gfp* into *osm-3*(*p802)* mutants. The two resulting independent lines, *jxEx194* and *jxEx195*, were also crossed to *daf-2*(*e1368*) to generate the *daf-2; osm-3* double mutants, in which *osm-3* was rescued from the ASJ neurons. To determine the effects of the *ins-6* UTRs on the *ins-6* mRNA subcellular localization, we created the lines *jxEx196* and *jxEx197* by injecting 2 ng/μl of pQZ97 with 25 ng/μl of the coinjection marker *ofm-1::gfp* into *ins-6*(*tm2416)* mutants. These were compared to the full *ins-6* rescuing lines *jxEx27* and *jxEx28*, which were injected with 2 ng/μl of pQZ11 and 25 ng/μl of the coinjection marker *ofm-1::gfp* into the *ins-6*(*tm2416)* mutants (*Cornils et al., 2011*). The coinjection marker *ofm-1::gfp* (25 ng/μl) has little or no effect on *ins-6* mRNA subcellular localization (*Figure 6B*) or dauer exit [*Table S1*; (*Fernandes de Abreu et al., 2014*)].

To test the sufficiency of the *ins-6* UTRs in trafficking *ins-6* mRNA to the axons, we injected 50 ng/μl of pQZ91 into the wild-type background, generating the extrachromosomal line *jxEx190*. This was compared to the extrachromosomal line *jxEx187*, which has been injected with 50 ng/μl of pQZ34 into the wild-type background and lacks the *ins-6* 3’ UTRs.

To confirm that dauer axons have Golgi bodies that can package newly synthesized INS-6 peptides, we injected 5 ng/μl of the pan-neuronal *rab-3p::aman-2::yfp* plasmid (generous gift of Stefan Eimer) with 25 ng/μl of the coinjection marker *ofm-1::gfp* into wild-type worms, generating the two lines *jxEx199* and *jxEx201*.

### Fluorescent *in situ* hybridization (FISH)

The mRNA of different ILPs or GFP were all visualized by FISH. Worms were washed with sterile water and fixed for 45 min with 4% paraformaldehyde at room temperature, ∼22°C (*Stewart et al., 2007*). After 3 five-min washes with phosphate-buffered saline (PBS), fixed worms were placed on a nutator and treated with 70% ethanol for 5 hrs to overnight at 4°C (*Tautz and Pfeifle, 1989*). Hybridization with the appropriate probe set (1:500 dilution) in hybridization buffer (15% deionized formamide, 10% dextran sulfate, 1 mg/ml *E. coli* tRNA, 4 mM vanadyl ribonuleoside complex and 0.2 mg/ml RNase-free BSA) was performed for 16 hrs in the dark at 30°C, followed by multiple 2-or 3-hr washes in 10% formamide in 2X SSC buffer for a total period of 36 hrs (*Raj and Tyagi, 2010*). The lower formamide concentration in the washes was to ensure retention of the signal during the washes. After the final wash, worms were kept at 4°C for 3 hours in 2X SSC buffer before treatment with Prolong Diamond Antifade (ThermoFisher, cat # P36961, refractive index of 1.47) for at least 24 hrs at 4°C.

All probes are labeled with CAL-Fluor 610 at the 3’ end and were generated through the Stellaris RNA-FISH Probe Designer 2.0, using the highest specificity (non-specificity masking level 5) from LGC Biosearch Technologies (Petaluma, CA).

### Image analyses of hybridized worms

Within a week of hybridization and Antifade-treatment, worms were mounted on 1% agarose pads and imaged on a Nikon Eclipse Ti-E or Ni-U microscope (Nikon Instruments Inc, Tokyo, Japan). Images were captured with a CoolSNAP MYO or ES2 CCD camera (Photometrics, Arizona, USA). Fluorescence intensities were quantified through a built-in fluorescence quantification algorithm (NIS-Elements/Annotations and Measurements/Mean Average Fluorescence Intensity, Nikon Instruments Inc), after a region-of-interest (ROI) background subtraction. Exposure time was kept constant at 900 ms without any neutral-density filter for all fluorescence imaging studies. All FISH images were taken at 400X magnification using the Nikon 40X N2 oil objective.

### Confocal laser scanning microscopy and live-image analyses

For live imaging of AMAN-2::YFP, well-fed L3 worms were mounted on 1% agarose pads with 5 mM sodium azide (Sigma-Aldrich, CAS No. 26628-22-8), whereas dauers were mounted on pads with 9 mM sodium azide (*Simpkin and Coles, 1981*). All confocal images were taken at 400X magnification within 30-40 min of mounting the worms.

#### Image acquisition

All confocal images and movies were acquired through a Leica laser-scanning confocal system (SP8), using a diode laser of 488 nm at 5% power (with a selected gated band-width of 500-560 nm, according to the Dye-Assistant plug-in of the LAS-X software). To achieve maximum sensitivity, emitted signals are detected through a photomultiplier tube at 500 nm. To obtain high signal-to-noise ratio from specimens, high-definition (1024 × 1024 dpi) images were scanned at a speed of 400 Hz/sec, with a line averaging of 4. To achieve true confocality with minimum photo-bleaching, the pinhole aperture was set to 1 airy unit of the 40X oil objective with a smart-gain value of 630.

To classify animals according to the amount of Golgi present in their dorsal cord axons, we used a particle-analyzer provided by FIJI (*Schindelin et al., 2012*). Animals with low axonal Golgi have less than 20 particles in a 50-μm section of the dorsal cord; animals with a medium amount of axonal Golgi have 21-40 particles in a 50-μm section of the dorsal cord; and animals with high axonal Golgi have greater than 40 particles in a 50-μm section of the dorsal cord.

#### Movie acquisition and 3D reconstruction

Using the protocol described above for image acquisition, each animal was scanned from bottom to top at 0.27-μm intervals to produce a stack of images, which were then bound together at three frames per second to produce a movie. To generate a 3D reconstructed image of an entire L3 or dauer larva from a 16-bit movie, we used the Leica 3D reconstruction program (LAS-X software; threshold = 20-120), which eliminated noise and puncta clusters (*Sibarita, 2005*). We used a volume-rendering method at 100% opacity during the 3D reconstruction of each animal (*Pawley, 2006*).

### Dauer exit assays

Dauer exit assays were performed according to *Cornils et al.* (*2011*). Eggs laid for 3-7 hrs at 20°C were collected and shifted to 25°C to induce dauer arrest. Forty-eight hours after the egg-laying midpoint, plates were checked for dauer formation, which were then assessed daily to calculate the rate of dauer exit.

### Statistical Analyses

All statistical analyses were performed using GraphPad Prism 7.0, except for dauer exit analyses. Wild-type *ins-6* hybridization signals were analyzed using 2-way ANOVA and Bonferroni correction with 95% confidence interval. Chi-square test was also performed to compare the differences between the classes of dauer populations across wild-type and mutant animals. For dauer exit, which were analyzed in JMP 6 from SAS, Kaplan-Maier probabilities were calculated and the *P-*value estimates were based on logrank comparisons. *P* < 0.05 was considered significant for all statistical comparisons.

## ACKNOWLEDGEMENTS

We thank S. Mitani, the *Caenorhabditis* Genetics Center (funded by NIH P40 OD010440), S. Eimer, A. Fire, M. Goodman, and P. Sengupta for strains and molecular reagents used in this study. We also thank R. Lillich and R. Patten of Mager Scientific Inc, Dexter, MI, and D. DeSantis and L. Mayernik of MICR Facility of Wayne State University for advice on fluorescence microscopy and the confocal scanning system; and Alcedo lab members, M. Friedrich, D. Njus, and S. Todi for discussions of this study. This work was supported by the Thomas C. Rumble University Graduate Fellowship to R. C. and by Wayne State University and NIH (R01 GM108962) to J. A.

