## Supplementary material for "Axonal Transport Of An Insulin-Like Peptide Mrna Promotes Stress Recovery In *C. Elegans*": Table S1

**Table S1. Kinesins that modulate axonal *ins-6* mRNA levels also modulate dauer exit**

| Strain | No. of animals observed/total animals | No. of trials | *P* vs. *daf-2(e1368)* | *P* vs. *daf-2(e1368)*; *osm-3*; *jxEx18* |
|---|---|---|---|---|
| **Kinesins that promote axonal *ins-6* mRNA** | | | | |
| *daf-2(e1368)* | 481/604 | 5 | - | na |
| *daf-2(e1368) unc-116* | 329/483 | 3 | < 0.0001 | na |
| *daf-2(e1368)*; *osm-3* | 52/588 | 3 | < 0.0001 | na |
| **Kinesin that inhibit axonal *ins-6* mRNA** | | | | |
| *daf-2(e1368)* | 258/347 | 3 | - | na |
| *daf-2(e1368) klp-6* | 312/381 | 3 | < 0.0001 | na |
| **ASJ-specific rescue of *osm-3* kinesin** | | | | |
| *daf-2(e1368)*; *jxEx18* | 193/229 | 2 | na | < 0.0001 |
| *daf-2(e1368)*; *osm-3*; *jxEx18* | 20/220 | 2 | < 0.0001 | - |
| *daf-2(e1368)*; *osm-3* | 10/114 | 2 | < 0.0001 | ns |
| *daf-2(e1368)*; *osm-3*; *jxEx194* | 64/135 | 2 | ns | < 0.0001 |
| *daf-2(e1368)*; *osm-3*; *jxEx195* | 82/163 | 2 | ns | < 0.0001 |

Statistical analyses are shown for the cumulative experiments in *Figure 5* of the dauer exit rates of *daf 2(e1368)* mutants in the presence or absence of specific kinesins at 25$^{o}$C. The presence of the coinjection marker *ofm-1::gfp*, *jxEx18*, has little or no effect on dauer exit, according to the statistical comparison between *daf-2*; *osm-3*; *jxEx18* and *daf-2*; *osm-3.* Many of the *daf-2*; *osm-3* double mutants, with or without the coinjection marker, crawl off the plates during the assay. However, the ASJ-specific rescue of *osm-3* partly rescues this “crawl-off” phenotype. To determine the statistical significance of the differences between groups, the logrank test is used. The following indicate: na, not applicable; ns, not significant, since $P > 0.05$.
