## Supplementary material for "Axonal Transport Of An Insulin-Like Peptide Mrna Promotes Stress Recovery In *C. Elegans*": Figure S1

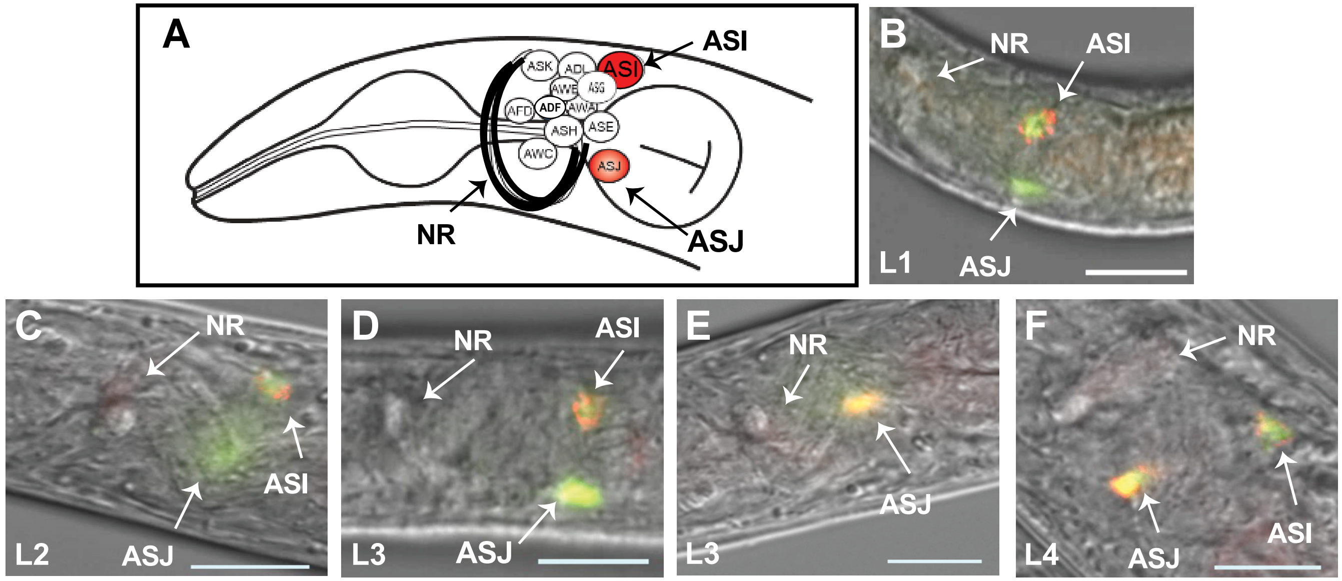


G

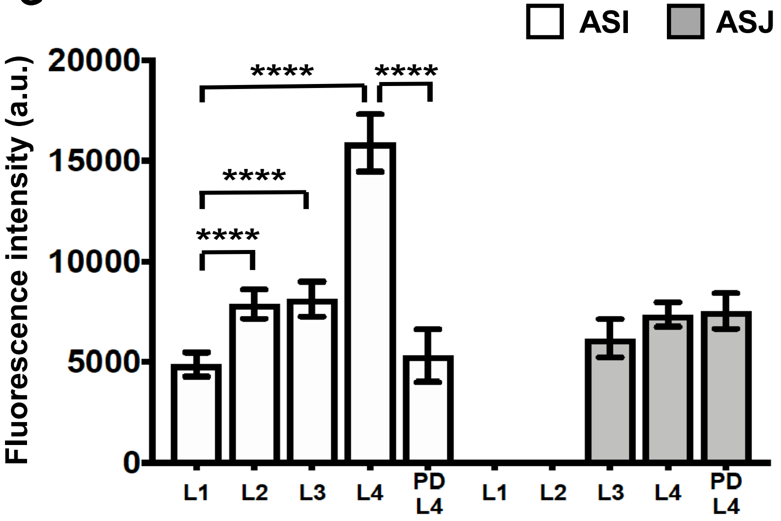


H

| stages | # of animals with neuronal *ins-6* mRNA | |
|---|---|---|
| | ASI | ASJ |
| L1 | 23/23 | 0/23 |
| L2 | 33/33 | 0/33 |
| L3 | 32/32 | 24/24 |
| L4 | 38/38 | 34/34 |
| PDL4 | 8/14 | 14/14 |

**Fig S1. *ins-6* mRNA is expressed in the larval ASI and/or ASJ sensory neurons during reproductive growth.** (A) Diagram of the 12 sensory neurons in the *C. elegans* amphid sensory organ (*White et al., 1986*). The ASI and ASJ sensory neurons are highlighted in red and NR indicates the nerve ring axon bundle. (B-F) An overlay of *ins-6* mRNA (red), GFP protein (green), and its respective DIC image in the same z-plane at 20°C in L1 (B), L2 (C), L3 (D-E) and L4 (F). Yellow represents colocalization of *ins-6* mRNA and the GFP protein expressed specifically in ASI and ASJ [*daf-28p::gfp*; (*Li et al., 2003*)]. In this figure and later figures, the anterior of each animal is to the left of each panel. Scale bar is 10 µm. (G) Mean fluorescence intensities (± SEM) of *ins-6* mRNA in ASI vs ASJ are shown from 2 trials for larval stages L1-L4 and post-dauer L4 (PDL4) in *daf-28p::gfp* animals (*Li et al., 2003*). The following indicates: a.u., arbitrary units; ****, $P < 0.0001$. (H) Distribution of larvae that express *ins-6* mRNA in ASI and/or ASJ in *daf-28p::gfp* animals from 2 trials.
