## Supplementary material for "Axonal Transport Of An Insulin-Like Peptide Mrna Promotes Stress Recovery In *C. Elegans*": Figure S2

**A**

**DAUERS**

| strains | # of animals with neuronal *ins-6* mRNA | | | mean total *ins-6* ± SEM | *P* vs *ins-6* | *P* vs ASJ-killed |
|---|---|---|---|---|---|---|
| | ASI | ASJ | NR | | | |
| wt A | 0/35 | 35/35 | 35/35 | 12427 ± 660 | < 0.0001 | < 0.0001 |
| wt B | 0/36 | 14/36 | 36/36 | 14851 ± 1452 | < 0.0001 | < 0.0001 |
| *ins-6* Δ | 0/36 | 0/36 | 0/36 | 1841 ± 225 | na | ns |
| ASJ-killed | 0/36 | 0/36 | 0/36 | 2515 ± 251 | ns | na |

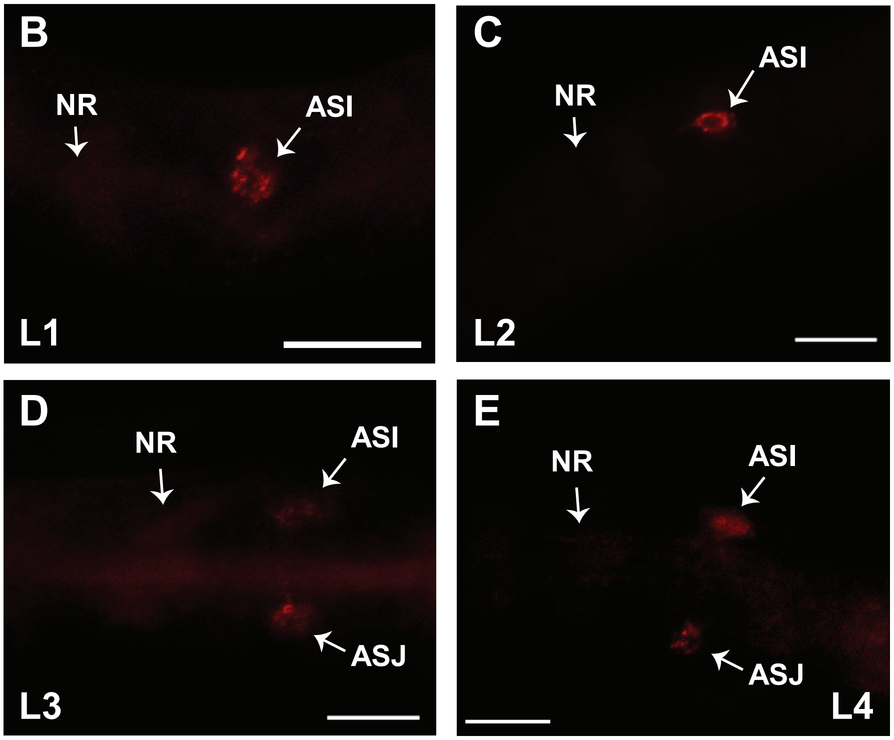


**Fig S2. *ins-6* mRNA is trafficked to the nerve ring (NR) axon bundle in starvation-induced dauers, but not in response to starvation alone.** (A) Distribution of wild-type (wt) or mutant dauers that express *ins-6* mRNA in ASI or ASJ soma or NR and mean total intensities (± SEM) of *ins-6* mRNA from 3 trials. (B-E) Representative fluorescent images of *ins-6* mRNA in starved wild-type L1 (B), L2 (C), L3 (D) and L4 (E) at 20°C. Animals were starved for about 5 days. Scale bar, 10 μm.
