## Supplementary material for "Axonal Transport Of An Insulin-Like Peptide Mrna Promotes Stress Recovery In *C. Elegans*": Figure S3

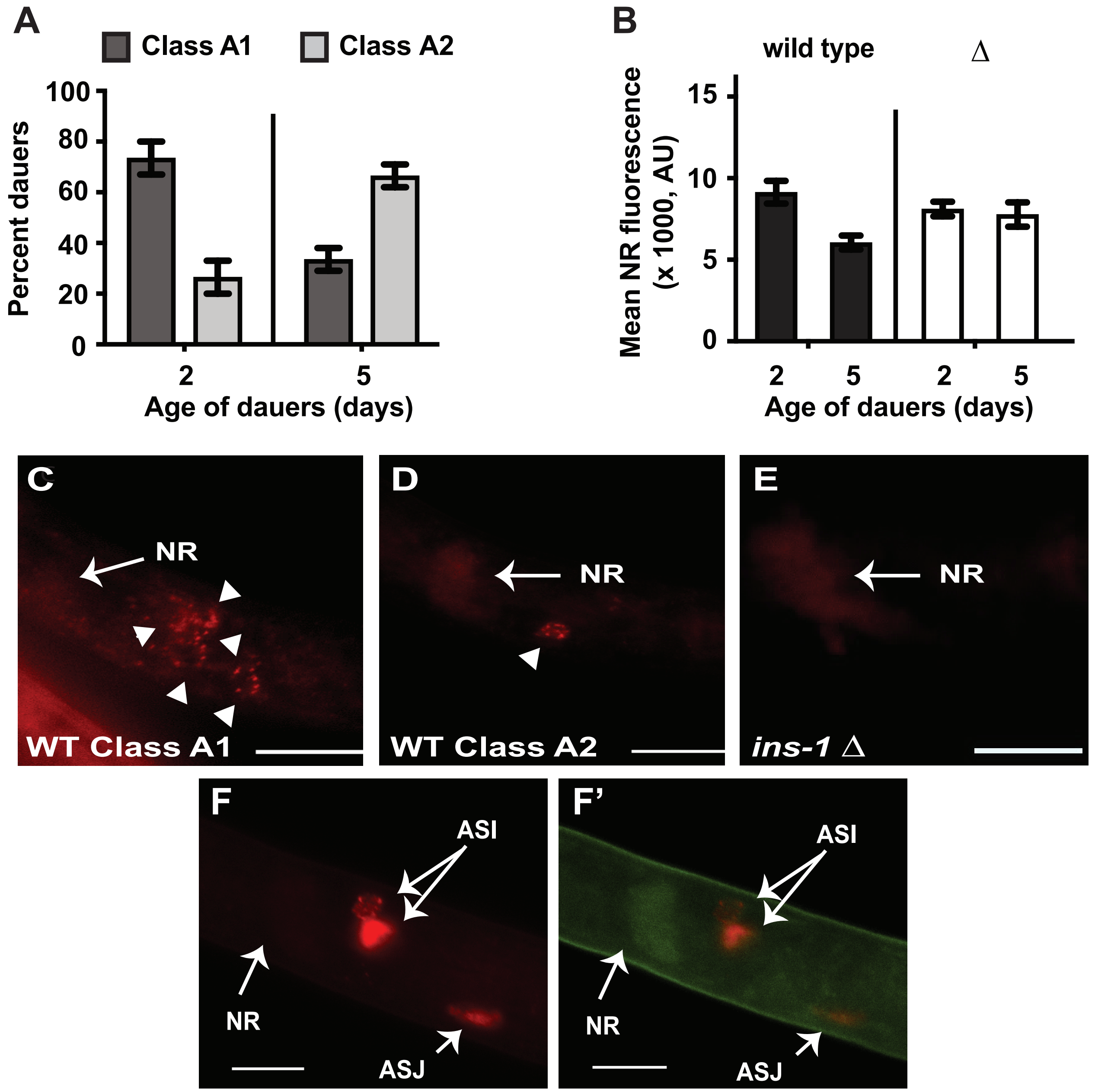


**Fig S3. Not all mRNAs are trafficked to the dauer NR axon bundle.** (A) Mean distribution (± SEM) of 2 classes of 2-day (n = 30) or 5-day (n = 36) old wild-type dauers at 20°C, based on number of neuronal somas that express *ins-1* mRNA. Class A1 dauers express *ins-1* mRNA in more neurons; class A2 dauers express *ins-1* mRNA in only one neuron. Values represent data from 4 trials. (B) Mean NR signals (± SEM) of *ins-1* mRNA in wild-type (from panel A) or *ins-1(nr2091)* deletion (Δ) mutant dauers. Number of *ins-1(Δ)* mutant dauers assayed for *ins-1* mRNA are 31 (2-day) and 29 (5-day) from 3 trials. AU, arbitrary units. (C-E) Fluorescent images of *ins-1* mRNA of 5-day old wild-type (C-D) and *ins-1(nr2091)* mutant (E) dauers at 20°C. Arrowheads denote the somas that express *ins-1* mRNA in wild-type dauers. (A-E) *ins-1* mRNA is assayed using probes specific to sequences deleted in the *ins-1(nr2091)* mutant. The NR shows similar background signals in both wild-type and *ins-1(Δ)* mutant dauers. (F-F') A 5-day old *daf-28p::gfp* dauer; F shows *gfp* mRNA (red); and F' is the merged image of *gfp* mRNA and protein (red and green, respectively). Scale bar, 10 μm.
