## Supplementary material for "Axonal Transport Of An Insulin-Like Peptide Mrna Promotes Stress Recovery In *C. Elegans*": Figure S4

**A**

| | Strains | *P* vs wild type |
|---|---|---|
| | wild type n = 25 | - |
| *daf-2* | *e1368* n = 25 | < 0.001 |
| *daf-2* | *e1370* n = 25 | < 0.001 |

**B**

| Strains | *P* vs wild type | *P* vs *klp-6* |
|---|---|---|
| wt n = 30 | - | < 0.001 |
| *unc-116* n = 32 | < 0.001 | < 0.001 |
| *osm-3* n = 16 | < 0.001 | < 0.001 |
| *klp-6* n = 26 | < 0.001 | - |

**C**

| | Strains | *P* vs wild type | *P* vs *osm-3* |
|---|---|---|---|
| | wt n = 20 | - | < 0.001 |
| | *osm-3* n = 25 | < 0.001 | - |
| *osm-3* | *jxEx194* n = 20 | ns | < 0.001 |
| *osm-3* | *jxEx195* n = 27 | ns | < 0.05 |

**D**

| | | | | | | | *e1368; osm-3* | |
|---|---|---|---|---|---|---|---|---|
| Strains | wild type n = 20 | *e1368* n = 25 | *osm-3* n = 20 | *klp-6* n = 25 | *e1368 klp-6* n = 20 | *e1368; osm-3* n= 38 | *jxEx194* n = 27 | *jxEx195* n = 28 |
| *P* vs wild type | - | < 0.0001 | < 0.0001 | < 0.01 | ns | < 0.0001 | ns | ns |
| *P* vs *e1368* | < 0.0001 | - | ns | < 0.0001 | < 0.05 | ns | ns | ns |
| *P* vs *osm-3* | < 0.0001 | ns | - | < 0.0001 | < 0.0001 | ns | < 0.001 | < 0.001 |
| *P* vs *e1368; osm-3* | < 0.0001 | ns | ns | < 0.0001 | < 0.0001 | - | < 0.0001 | < 0.0001 |

**Fig S4. Insulin signaling and specific kinesins regulate trafficking of *ins-6* mRNA to dauer axons.** (A-D) Statistical comparisons across different strains from the corresponding panels in *Figure 4*. Statistical analyses are determined by two-way ANOVA and Bonferroni correction. Values represent data from at least 2 trials.
